# FtsK with a unique N-terminal extension is involved in coordinating the final steps of chromosome segregation with asymmetrical division in mycobacterial cells

**DOI:** 10.64898/2026.01.27.701935

**Authors:** Kornel Milcarz, Joanna Hołówka, Damian Trojanowski, Tomasz Łebkowski, Dominik Bania, Michał Tracz, Jolanta Zakrzewska-Czerwińska

**Affiliations:** Department of Molecular Microbiology, University of Wrocław, Wrocław, Poland; Protein Mass Spectrometry Laboratory, University of Wrocław, Wrocław, Poland

## Abstract

In this study, we identify and functionally characterize a previously unrecognized, conserved N-terminal extension of mycobacterial FtsK (nFtsK) that contributes to the coordination of chromosome segregation with asymmetric cell division. Although FtsK is broadly conserved in bacteria as a late-stage DNA translocase and divisome component, mycobacterial FtsK uniquely contains a long, positively charged, intrinsically disordered N-terminal region. We show that this extension is dispensable for core FtsK functions—including septal localization, DNA translocation, and completion of cell division—but instead modulates the spatial confinement, temporal regulation, and interaction specificity of these processes. Mechanistically, nFtsK mediates specific protein interaction, binds the anionic phospholipids cardiolipin and phosphatidic acid, and exhibits intrinsic affinity for negatively curved membrane. Together, our findings uncover a lineage-specific adaptation that fine-tunes a conserved molecular machine, enhancing the robustness of asymmetric cell division in mycobacteria.

## Introduction

Most rod-shaped bacteria grow by lateral elongation and divide symmetrically via septum formation at mid-cell, producing two daughter cells of equal lengths, each inheriting a complete copy of genome. These processes have been extensively studied in model organisms such as the Gram-negative *Escherichia coli* and the Gram-positive *Bacillus subtilis*. In these species, symmetric division is tightly regulated by spatial control systems—including the MinCDE and nucleoid occlusion (NO) systems—that ensure accurate placement of the FtsZ ring (Lutkenhaus, 2007; Wu & Errington, 2012). These mechanisms prevent mispositioning of the division machinery and protect against lethal guillotining of the chromosome(s).

In contrast, *Mycobacterium* species exhibit a markedly different mode of growth and division. These bacteria elongate in a polar, biphasic manner, with the old pole growing more rapidly than the newly formed pole, and divide asymmetrically, generating daughter cells of unequal lengths and different physiology (Aldridge et al., 2012; Kieser & Rubin, 2014). Notably, no homologs of the MinCDE or nucleoid occlusion (NO) systems have been identified in mycobacteria to date, raising fundamental questions about how chromosome segregation and cell division are coordinated to preserve genome integrity.

The early steps of chromosome segregation in mycobacteria are relatively well characterized. The ParAB*S* system plays a central role in positioning newly replicated *oriC* regions toward the cell poles: ParB binds centromere-like *parS* sequences near *oriC* to form nucleoprotein complexes (segrosomes), which are actively segregated by the ATPase ParA (Ginda et al., 2017; Jakimowicz et al., 2007). However, the later steps of segregation – particularly DNA translocation across the closing septum – remain poorly understood.

In most bacteria, late-stage chromosome segregation is mediated by FtsK, an essential DNA translocase and core divisome component (Bush et al., 2025; Cameron & Margolin, 2024). In *E. coli*, FtsK localizes to the septum during late-cytokinesis, and deletion of its C-terminal motor domain results in defects in both chromosome segregation and septation. FtsK assembles into a hexameric, ATP-driven motor that actively translocates double-stranded DNA from the division site into the nascent daughter cell (Jean et al., 2020). Its C-terminal domain comprises three subdomains: α and β, which form the hexameric motor, and γ, which controls directionality and interacts with the XerCD recombinase complex (Yates et al., 2006). FtsK recognizes G-rich KOPS (FtsK orienting polar sequences) motifs that guide DNA translocation toward the *dif* site near the replication terminus (*ter*) region. These motifs are asymmetrically distributed along each chromosome arm, oriented toward *dif* and progressively enriched. Translocation terminates through interaction with the XerCD recombinase complex, facilitating chromosome dimer resolution and decatenation (Sivanathan et al., 2009). Beyond translocase activity, the N-terminal domain of FtsK anchors the protein to the cytoplasmic membrane via transmembrane helices and mediates interactions with other divisome components (e.g., FtsQ, FtsL, FtsI), thereby coupling cell division with chromosome segregation (Grenga et al., 2008). Studies in other organisms (e.g., *Staphylococcus aureus*, *Deinococcus radiodurans*) further demonstrate that FtsK indirectly influences the levels of the septal peptidoglycan hydrolase Sle1, divisome assembly, and the timing of septal splitting, underscoring its dual role in DNA translocation and coordination of cell division (Veiga et al., 2023).

In mycobacteria, FtsK is essential (DeJesus et al., 2017; Griffin et al., 2011; Minato et al., 2019; Sassetti et al., 2003) but it remains largely uncharacterized. Notably, mycobacterial FtsK contains a uniquely extended, intrinsically disordered, and positively charged N-terminal region (see **Fig. 1A**), suggesting a lineage-specific adaptation linked to polar growth and asymmetric division. Here, we show that this conserved N-terminal region enhances the robustness and precision of late chromosome segregation by constraining translocase dynamics and septal positioning. Loss of this region does not abolish FtsK function but leads to faster yet less precisely coordinated cell cycle progression.

**Figure 1.**
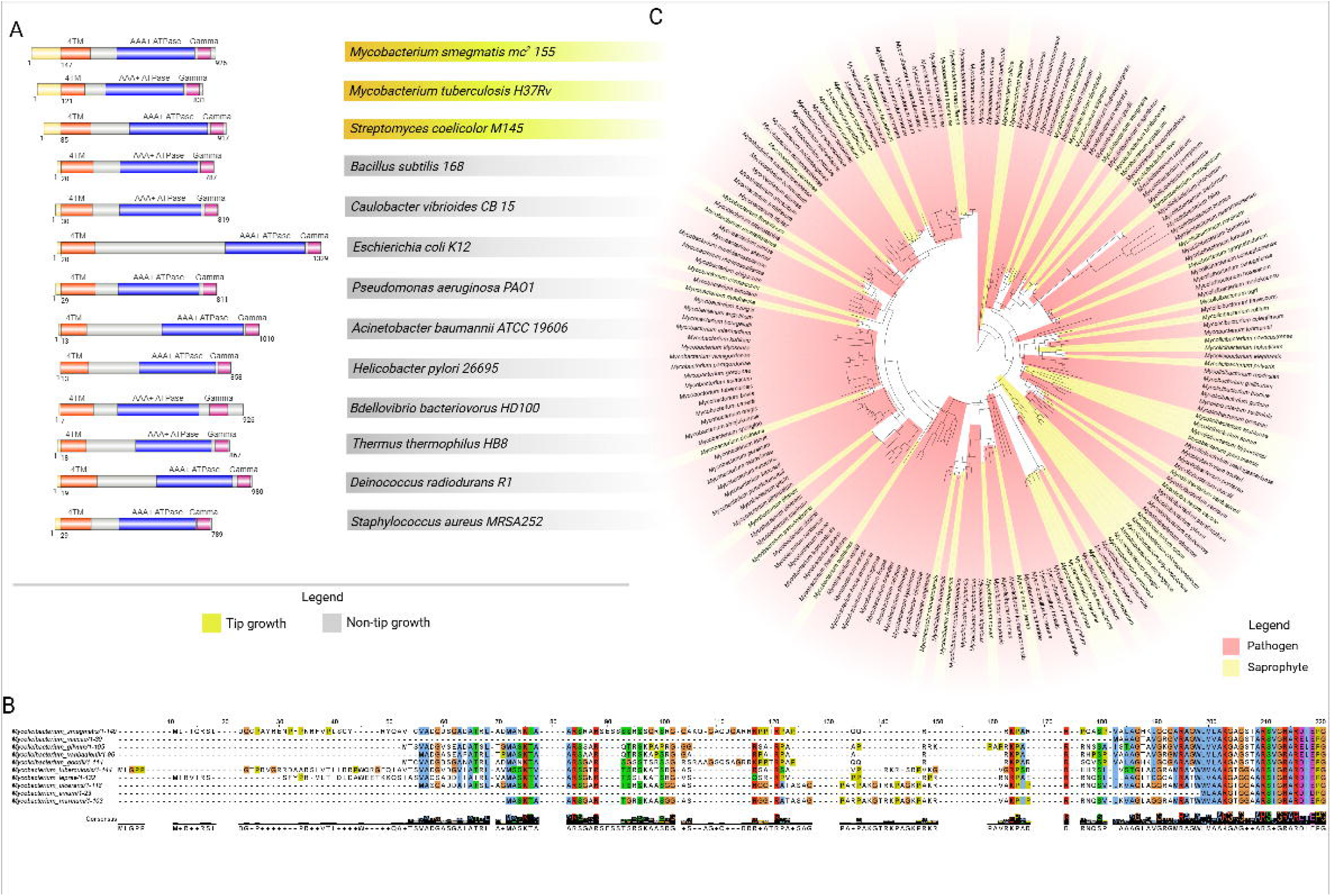
Comparative analysis and conservation of FtsK across diverse bacterial species and within *Mycobacterium* genus. **A.** Comparison of FtsK domain organization in *Mycobacterium* species and representative bacterial model organisms. The N-terminal extension is indicated in yellow, followed by four transmembrane helices (4TM) shown in orange. The linker region is depicted in grey, while the AAA+ ATPase motor domain and the gamma domain, responsible for KOPS interaction, are shown in blue and violet, respectively. **B.** Multiple sequence alignment of FtsK proteins from selected bacterial species and representative *Mycobacterium* strains. Shown is an enlarge view of the N-terminal 150 amino acids present in *M. smegmatis*; the full alignment is provided in the Supplementary Materials. **C.** Phylogenetic tree of FtsK sequences from 150 *Mycobacterium* species.

## Results

### *In silico* analysis reveals a conserved N-terminal extension in mycobacterial FtsK

Variation among FtsK proteins is primarily observed in their N terminal domains, which typically contain four transmembrane helices and a linker connecting the membrane anchor to the well conserved cytosolic translocation motor (Crozat & Grainge, 2010). To assess whether mycobacterial FtsK differs from homologs in other bacteria, we compared amino acid sequences from representative species. Sequence alignment revealed that mycobacterial FtsK possesses a unique, extended N-terminal tail (**Fig. 1A**). In most Gram-positive and Gram-negative bacteria, the first transmembrane helix begins immediately after the initiator methionine. In contrast, mycobacterial FtsK contains an additional >150 amino acids upstream of these helices (**Fig. 1A**). A similar but shorter extension (approximately 80 amino acids) was observed in *Streptomyces coelicolor*, another member of the phylum *Actinomycetota*.

This N-terminal extension (here and after referenced as nFtsK) is conserved among mycobacterial species, with slight differences in its length. Alignment of ten representative saprophytic and pathogenic species revealed a conservation embedded within a nFtsK fragment (**Fig. 1B, lower panel**). In *M. smegmatis* this extension is strongly positively charged (net charge +23, pI = 12) and hydrophilic, and enriched in Arg (15%), Lys (3.9%), Ser (10%), and Gly (10%), suggesting that this segment is intrinsically disordered. Phylogenetic analysis further showed that FtsK sequences do not cluster according to saprophytic or pathogenic lifestyle, indicating that this extension represents a conserved evolutionary adaptation across the genus.

In summary, mycobacterial FtsK contains a conserved, positively charged and hydrophilic N-terminal extension that is absent from most other bacteria lineages.

### Truncation of the N-terminal extensions alters FtsK dynamics and accelerates the cell cycle

Given the lack of lifestyle specificity of nFtsK and the high conservation of this region among mycobacteria (**Fig. 1C**), we next investigated its role. We initially attempted to delete 5’-endfragment of the *ftsK* gene encoding the entire unstructured N-terminal region. However, no viable double-crossover (DCO) strains were obtained, suggesting that a 4 transmembrane _α_-helix domain is required for FtsK function. We therefore constructed a truncated variant lacking only the first 140 amino acids while preserving the first predicted α-helix. This variant, termed sFtsK (short FtsK), was fused to a HaloTag (sFtsK-HT) and compared with the full-length protein FtsK-HT.

The sFtsK-HT strain exhibited normal morphology but displayed a modest yet significant reduction in mean cell length compared with the wild type (3.75 ± 1.00 µm vs. 4.21 ± 1.01 µm; p = 1.6 × 10^-3^, n = 104). Both strains exhibited comparable growth profiles; however, a slight, non-significant reduction in growth rate was observed for the sFtsK-HT strain relative to the wild-type strain. FtsK-HT and sFtsK-HT formed distinct signal at the septum (**Fig. 2A**) and colocalized with established septal markers (**Fig. 2B**), indicating that the N-terminal extension is not required for septal recruitment. Moreover, 3D-SIM analysis showed that FtsK-HT shapes into ring-like structure (**Movie 1**). Interestingly, the population-level analysis revealed that sFtsK-HT localized slightly closer to midcell compared with the full-length protein (**Fig. 2C**), suggesting reduced spatial confinement in the absence of the N-terminal extension of FtsK. Despite similar recruitment dynamics, the residence time of FtsK at the septum was significantly reduced upon removal of the N-terminal extension. Full-length FtsK-HT persisted at the septum for 62 ± 24 min, whereas sFtsK-HT remained for only 50 ± 14 min (p = 2.23 × 10^-8^; n = 216 and 189, respectively) (**Fig. 2D, top right**).

**Figure 2.**
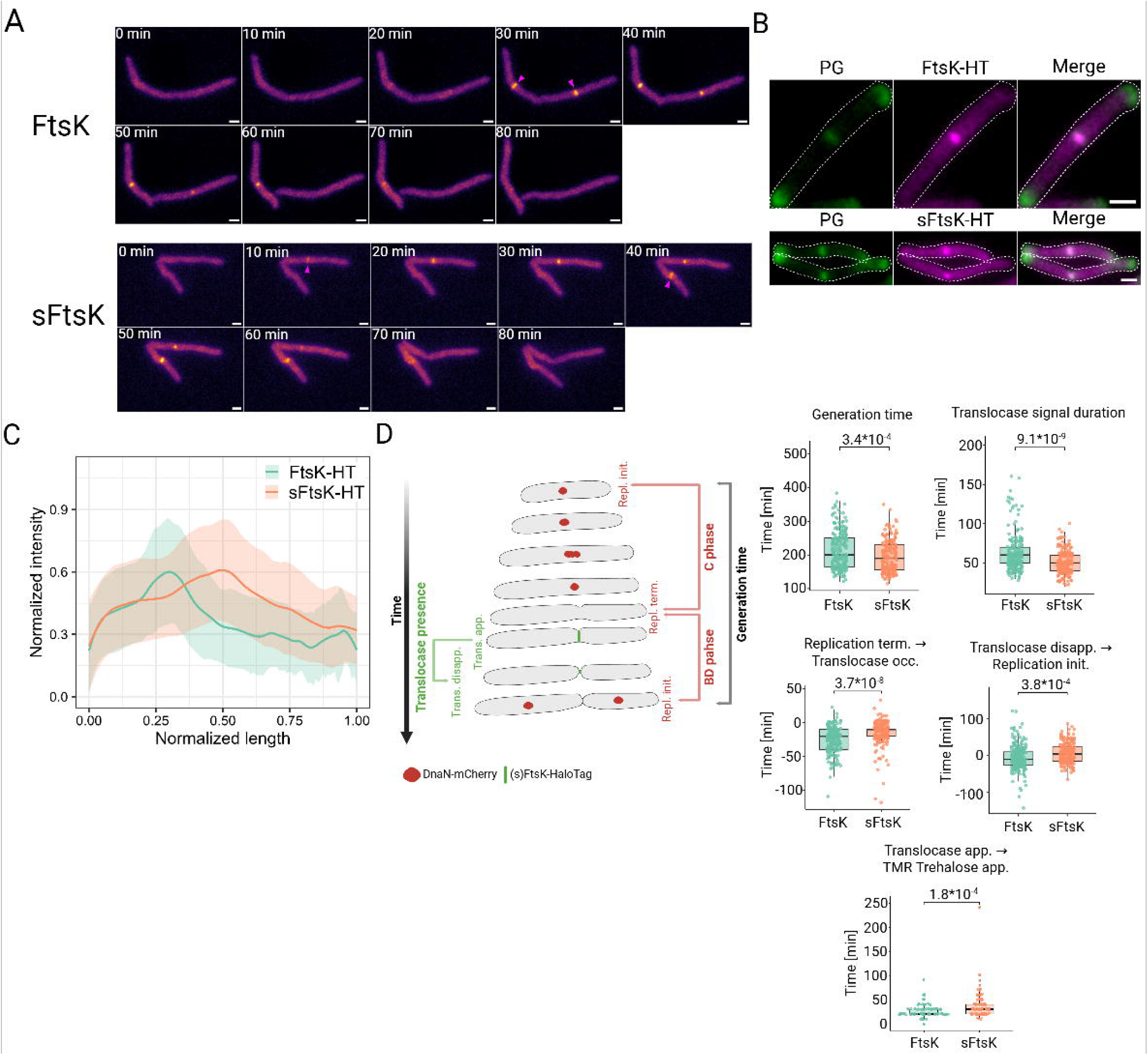
The truncated sFtsK variant exhibits altered localization and accelerated cell cycle dynamics. **A.** Representative time-lapse fluorescence microscopy images of *M. smegmatis* cells expressing FtsK-HT (top panels) or the truncated sFtsK-HT variant (bottom panels). Magenta arrowheads indicate appearance of representative translocase foci. Scale bar, 1 µm. **B.** Representative images showing colocalization of sFtsK-HT with the peptidoglycan marker NADA-Green, scale bar, 1 µm. **C.** Population-level profiles of normalized sFtsK-HT and FtsK-HT fluorescence intensity plotted against normalized cell length, showing that loss of nFtsK shifts the position of the fluorescence intensity maximum (n = 197 and 151 for FtsK-HT and sFtsK-HT, respectively). **D.** Left: schematic of the cell cycle highlighting the dynamics of DNA replication and sFtsK localization on left panel. Right: quantification of cell cycle parameters and translocase dynamics in FtsK-HT (teal) and sFtsK-HT (orange) strains. Plots show generation time (top left), translocase signal duration (top right), the time interval between replication termination and translocase appearance (bottom left), the time interval between translocase disappearance and subsequent replication initiation (bottom right) and the time interval between translocase appearance and TMR trehalose appearance (cell envelope marker). Each dot represents a single cell.

To place these localization differences in the context of the cell cycle, we monitored DNA replication using the replisome marker DnaN-mCherry (Trojanowski et al., 2017), which allowed tracking of the C phase (replication duration) and the BD phase (the interval between replication termination and initiation of the next round in daughter cells). Generation time, defined as the interval between replication focus appearance in mother and daughter cells, was significantly shorter in sFtsK-HT-expressing cells that in FtsK-HT cells (195 ± 47 min vs. 214 ± 56 min; p = 3.96 × 10^-4^; n = 189 and 216, respectively) (**Figure 2D, top left**), consistent with the reduced duration of sFtsK septal localization. Relative to replication termination, FtsK-HT foci appeared 26 ± 18 min before termination, whereas sFtsK-HT foci emerged only 15 ± 18 min earlier (p = 3.62 × 10LL, n = 216 and 189, for FtsK-HT and sFtsK-HT respectively) (**Fig. 2D, bottom left**). In daughter cells, replication initiation occurred later in the sFtsK-HT strain, whereas in FtsK-HT cells it tended to begin slightly before translocase disappearance (6 ± 27 min vs −5 ± 34 min; p = 4.63 × 10LL, n = 189 and 216, for sFtsK-HT and FtsK-HT, respectively) (**Fig. 2D, bottom right**). Notably, both strains exhibited substantial heterogeneity, reflecting the flexible coupling between DNA replication and cytokinesis in *Mycobacterium*. We next assessed whether this flexibility extended to cell envelope biogenesis by measuring septal TMR-trehalose incorporation (**Fig. 2D**, **bottom center**). While both strains showed similar modes of signal appearance, incorporation was significantly delayed and more heterogeneous in sFtsK-HT cells compared with the full-length FtsK cells (38 ± 26 min vs 27 ± 12 min; p = 1.80*10^-4^, n = 110 and 107, respectively). The increased variance in sFtsK-HT cells is consistent with impaired temporal coordination between septal cell wall synthesis and the cell cycle following removal of the N-terminal extension.

Given these temporal shifts, we next examined whether coordination between translocation termination and chromosome segregation was preserved. Analysis using the nucleoid marker HupB revealed that the mean timing of chromosome segregation relative to translocase disassembly was indistinguishable between the two variants (p > 0.67). However, sFtsK-HT exhibited significantly greater variability—approximately 40% higher—in this coordination compared with full-length FtsK-HT (variance: 281 vs. 198; Levene’s test, p = 0.01). This increased stochasticity suggests that the extended N-terminal region is required for the precise temporal coupling of translocation termination with chromosome segregation, rather than for the overall rate of the process.

Together, these data demonstrate that while the N-terminal extension of mycobacterial FtsK is dispensable for septal recruitment and overall divisome function, it fine-tunes the spatial stability and temporal precision of FtsK activity, thereby enhancing robust coordination between cell division, DNA replication, and chromosome segregation.

### The N-terminal extension constrains FtsK diffusion and spatial confinement

To investigate the molecular basis of the altered cell cycle timing, we performed single-particle tracking (SPT) of FtsK-HT and the truncated variant sFtsK-HT. In total, 9951 trajectories were analyzed for FtsK-HT and 8843 for sFtsK-HT. Mean squared displacement (MSD) analysis revealed similar overall diffusion trends for both variants: MSD increased with the time lag (τ), but its rate of decreased over time, consistent with subdiffusive behavior (**Fig. 3A**). Corresponding with the reduced MSD, the average diffusion coefficient was lower for sFtsK-HT than for FtsK-HT (D_xy_ = 0.053 µm^2^/s vs. 0.078µm^2^/s; track/cell = 59 and 76, n = 169 and 134, respectively) (**Fig 3A**).

**Figure. 3.**
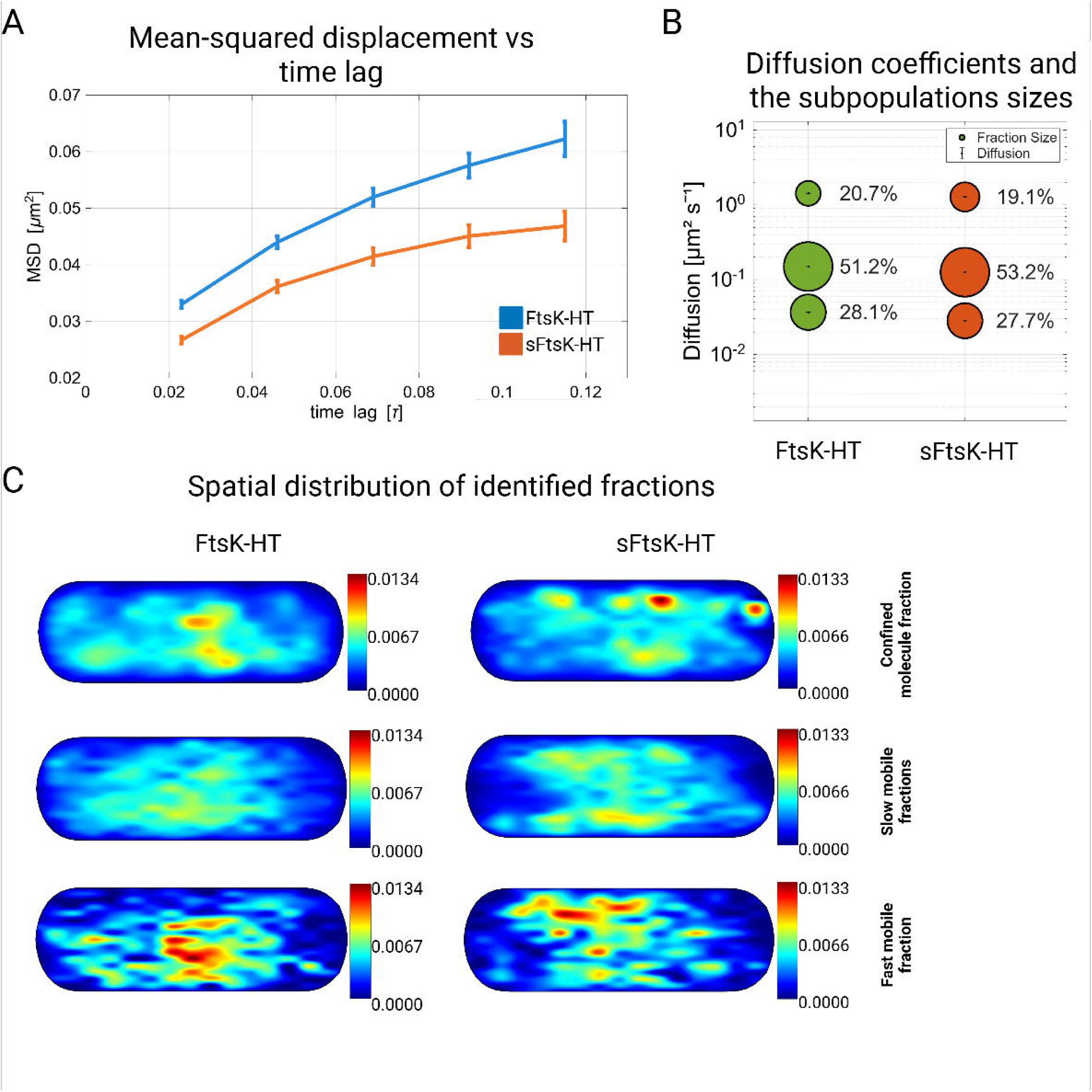
Single particle tracking analysis of FtsK-HT and sFtsK-HT. **A.** Mean Squared Displacement (MSD) plotted as a function of time lag. The sFtsK-HT variant (orange line) exhibits a lower slope than FtsK-HT (blue line), indicating reduced motion. **B.** Diffusion coefficients and subpopulation sizes. The bubble plot shows diffusion coefficient (y axis) and relative fraction size (bubble area and percentage) for three dynamic states: fast mobile, slow mobile, and confined molecules. A shift in population equilibrium is observed in the sFtsK strain, characterized by an overall reduction in diffusion coefficients. **C.** Spatial probability density maps of identified fractions. Heatmaps show the localization of particles belonging to the confined, slow mobile, and fast mobile populations projected onto a normalized cell contour. The color scale represents particle density ranging from low (blue) to high (red). The slow-mobile fraction of full-length FtsK-HT preferentially accumulates at mid-cell, whereas the sFtsK-HT variant displays a more dispersed localization pattern lacking central confinement.

To further resolve diffusive heterogeneity, we performed square displacement (SQD) analysis using a non-simultaneous parameter approach. Although the relative population sizes of diffusive states were similar between the two variants, their dynamic properties differed markedly (p = 8.67 × 10^-64^, **Fig. 3B**). Specifically, diffusion coefficients were consistently reduced for sFtsK-HT across all three mobility states. The fast mobile fraction slowed down from D_xy_ = 1.430 ± 0.003 µm^2^/s in FtsK-HT to 1.280 ± 0.003 µm^2^/s in sFtsK-HT. Likewise, both the slow mobile (0.150 vs. 0.126 µm^2^/s) and confined fractions (0.037 vs. 0.028 µm^2^/s) exhibited reduced mobility in the absence of the extended N-terminal region. Despite these changes in diffusion rates, the relative proportions of the three populations remained comparable between variants (from most mobile to least mobile: 20.7% vs. 19.1%, 51.2% vs. 53.2%, 28.1% vs. 27.7% for FtsK-HT and sFtsK-HT, respectively).

We next assessed immobilization kinetics using dwell time analysis with a 100-nm radius and a two-component model. The characteristic residence times for transient (τ_1_ ≈ 0.16s) and stable (τ_2_ ≈ 0.38s) dwell events were indistinguishable between the two variants. In contrast, removal of the extended N-terminal region significantly increased the probability of entering the stable immobilized state. Specifically, sFtsK-HT exhibited an approximately 40% relative increase in the stable dwell fraction (τ_2_), rising from 12.3% to 17.3% (p = 5.82 × 10^-11^). These results indicate that loss of the extended disordered N-terminal region promotes transitions into confined states without altering the intrinsic stability of immobilized state.

Consistent with these finding, spatial mapping of diffusion clusters revealed pronounced differences in localization patterns between the two variants (**Fig. 3C**). Full-length FtsK-HT exhibited robust mid-cell enrichment across all diffusive clusters, including the fastest population (D_xy_ = 3.49 µm^2^/s). In contrast, sFtsK-HT displayed a broader and more delocalized distribution, with only the intermediate mobility cluster (D_xy_ = 0.42 µm^2^/s) enriched at mid-cell. Moreover, the slowest diffusive clusters differed substantially between variants (D_xy_ = 0.124 for FtsK-HT vs. 0.012 µm^2^/s for sFtsK-HT). Notably, only full-length FtsK-HT exhibited a static, confined population, whereas sFtsK-HT retained Brownian-like motion even in its slowest state.

Collectively, these results demonstrate that the extended N-terminal region of FtsK is essential for promoting spatial clustering and constrained dynamics at the division site. Although removal of this region does not abolish septal localization, it shifts FtsK dynamics toward slower, more delocalized diffusion and increases promiscuous immobilization events. We propose that the extended N-terminal region fine-tunes FtsK confinement by suppressing off-target interactions, potentially with the membrane or non-divisome proteins, thereby ensuring high-fidelity localization at the divisome.

### The N-terminal region of FtsK mediates specific protein and lipid interactions

Single-particle tracking revealed that the removal of the extended N-terminal region of FtsK results in reduced molecular mobility and a loss of spatial confinement at the mid-cell, leading to a more dispersed localization pattern. Importantly, this defect did not abolish septal complex formation by sFtsK, indicating that the N-terminal region is not essential for core divisome recruitment. Instead, the altered dynamics suggested that the N-terminal region (nFtsK) may modulate interactions with ‘secondary’ partners or suppress non-specific associations.

Given the net positive charge of nFtsK, we first tested whether this region mediates DNA binding. However, electrophoretic mobility shift assay (EMSA) revealed no detectable interaction between purified nFtsK (**Fig. S2**) and DNA, as no protein–DNA complexes were observed under the condition tested (**Fig. 4A**). This data indicate that nFtsK does not function as an autonomous DNA-binding module.

**Figure 4.**
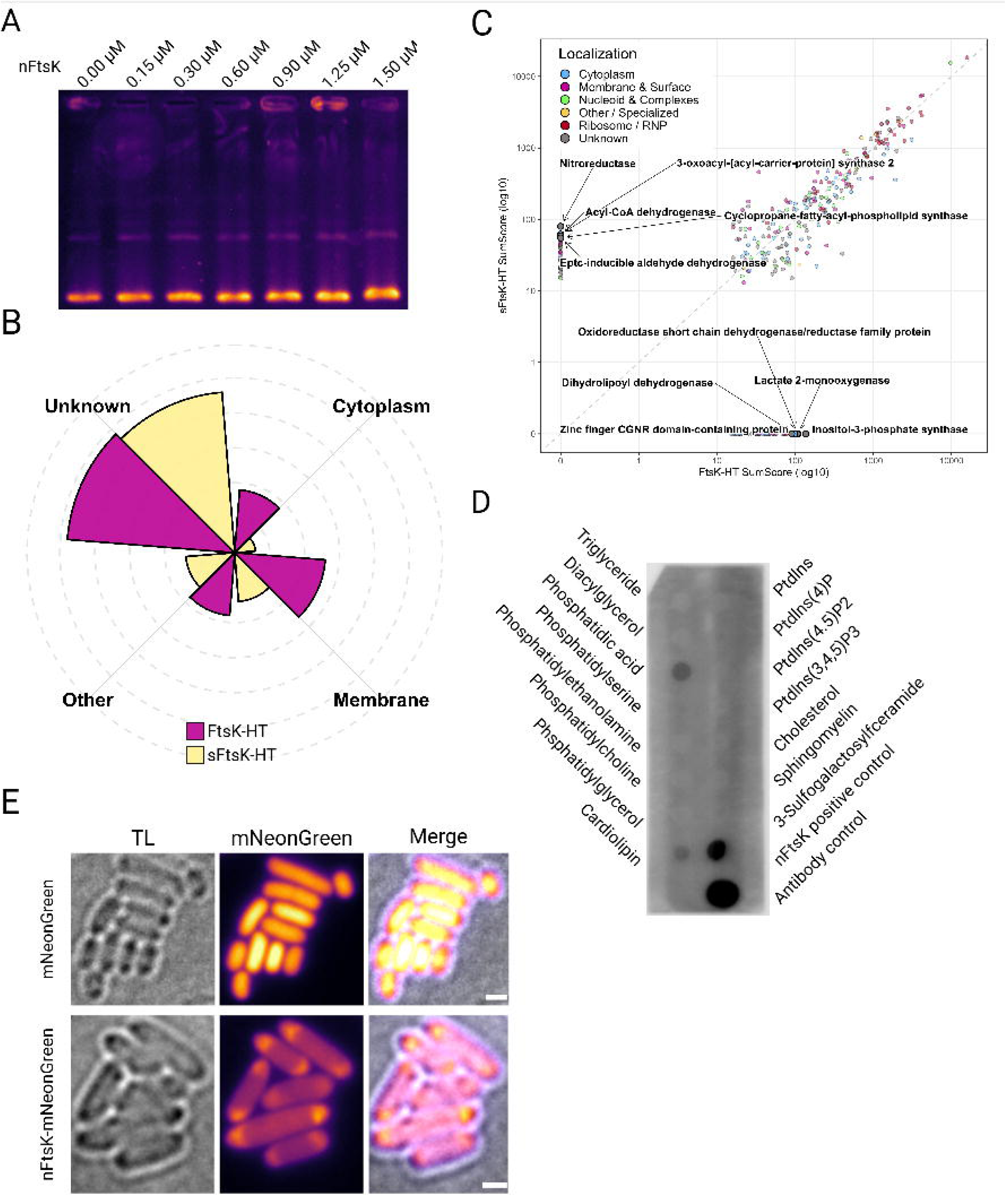
Functional characterization and interactome profiling of the N-terminal region of FtsK (nFtsK). **A.** Electrophoretic mobility shift assay (EMSA) assessing the DNA-binding activity of purified nFtsK, showing no protein-DNA interaction. **B.** Subcellular localization profiling of proteins identified exclusively in FtsK-HT or sFtsK-HT pulldowns. Functional categories were assigned based on UniProt “Subcellular location [CC]” annotations and transmembrane domain predictions. **C.** Comparative scatter plot of the FtsK-HT and sFtsK-HT interactomes. Axes represent log_10_ transformed enrichment intensities. Labeled points indicate the top unique interactors of the full-length (FtsK-HT) and truncated (sFtsK-HT) variants, highlighting distinct interaction landscapes. **D.** Lipid-protein overlay assay demonstrating the specificity of nFtsK for cardiolipin and phosphatidic acid. **E.** Representative image of nFtsK-mNeonGreen expressed in the heterologous host *E. coli*, showing preferential localization to membrane regions with negative curvature. Scale bar, 1 µm.

We next examined whether nFtsK contributes to specific protein-protein interactions by comparing the interactome of FstK-HT and sFtsK-HT (**Table S1**). Both variants co-purified robustly with established septal and divisome-associated proteins, including MSMEG_4287 (a recently identified septal factor), MSMEG_5223, SecA1, Ffh, and OxaA, the latter three being involved in membrane proteins targeting. FtsK itself, as well as numerous proteins previously identified in FtsQ pulldowns, a close divisome partner, were also detected (Wu et al., 2018). Importantly, SepF, a membrane anchor for the Z-ring, was present in both datasets. Additionally, the nucleoid-associated proteins HupB and mIHF, as well as the chaperons GroEL2 and GroES, were detected in both interactomes. Together, these findings indicate that truncation of the N-terminal region does not disrupt recruitment of core divisome components, consistent with the preserved septal localization observed for sFtsK.

Despite this overall similarity, comparative analysis revealed a distinct set of proteins that were selectively enriched in the FtsK-HT interactome, implicating nFtsK in additional regulatory interactions (**Fig. 4B and C**). Notably, Ino1, the rate-limiting enzyme of inositol biosynthesis, and a key contributor to phosphatidylinositol-derived cell envelope components (phosphatidylinositol mannosides PIMs, lipoarabinomannan LAM, and lipomannan LM) was detected exclusively in the FtsK-HT pulldown. Several enzymes involved in acetyl-CoA production, critical for mycolic acid and peptidoglycan synthesis, were also uniquely associated with FtsK-HT, including two components of the pyruvate dehydrogenase complex (PDH) and lactate 2-monooxygenase, which converts lactate to acetate, a direct precursor of acetyl-CoA. In addition, MSMEG_1813, required for lipid branching reactions, and WhiA, a known coordinator of cell division, were identified only in the presence of full-length FtsK. These associations suggest that nFtsK links septal translocation with metabolic pathways essential for cell envelope biogenesis.

In contrast, the sFtsK-HT interactome was characterized by increased enrichment of proteins indicative of aberrant or promiscuous interactions. The membrane protease FtsH, which targets misfolded or damaged membrane associated proteins, was markedly enriched in the sFtsK-HT pulldown, exhibiting 13-fold higher enrichment compared to FtsK-HT. This suggests that removal of the terminal region renders FtsK more susceptible to quality control surveillance. Additionally, the δ subunit of DNA Polymerase III (a replisome component) was detected exclusively in the sFtsK-HT fraction. Ribosomal proteins, commonly associated with non-specific binding in pulldown experiments, were also enriched in the truncated variant. These observations are consistent with the increased non-specific immobilization and delocalized diffusion detected for sFtsK *in vivo*.

Finally, we examined whether nFtsK interacts with membrane lipids. Lipid-binding assays revealed that nFtsK preferentially associates with phosphatidic acid (PA) and cardiolipin (CL), two canonical lipids that accumulate in regions of negative membrane curvature (Mileykovskaya et al., 2009; Renner & Weibel, 2011), such as the septum and cell poles (**Fig. 4D**). To determine whether this interaction is intrinsic to nFtsK and not mediated by co-purifying septal proteins, we performed *ex vivo* localization assays. Fusion of nFtsK to mNeonGreen was sufficient to direct the fluorescent reporter to cell poles, regions known to be enriched in CL and PA (**Fig. 4E**).

Collectively, these results demonstrate that the N-terminal region of FtsK engages in specific protein-protein and protein-lipid interactions. By associating with negatively curved membrane domains and metabolic regulators of cell envelope synthesis, nFtsK promotes spatial confinement and suppresses non-specific interactions, thereby stabilizing FtsK positioning and dynamics at the division site.

## Discussion

Unlike laterally growing, symmetrically dividing model bacteria, mycobacteria elongate from the cell poles and divide asymmetrically (Aldridge et al., 2012; Chung et al., 2024) in the absence of the canonical MinCDE and nucleoid occlusion systems that ensure accurate septum placement and protect the chromosome. Asymmetric division produces daughter cells with distinct sizes and growth capacities, generating phenotypic heterogeneity within the population (Aldridge et al., 2012; Chung et al., 2024). Such heterogeneity is thought to act as a bet-hedging strategy that promotes survival under stress, including antibiotic challenge (Aldridge et al., 2012). Despite the lack of these spatial regulators, septum formation and closure occur with high fidelity, implying the existence of alternative mechanisms that couple divisome activity to chromosome dynamics. This raises the fundamental question of how late-stage chromosome segregation is coordinated with septum constriction to prevent guillotining of the chromosome and ensure genome integrity.

In mycobacteria, the essential DNA translocase FtsK (DeJesus et al., 2017; Griffin et al., 2011; Minato et al., 2019; Sassetti et al., 2003) functions during the late stages of chromosome segregation, as in other bacteria. FtsK is recruited only after assembly of core divisome, coinciding with the onset of septal constriction, and localizes precisely to the leading edge of the invaginating septum, forming a ring-like structure that ensures DNA is cleared before final scission (**Fig. 2A, Movie 1**). Unlike its homologs in symmetrically dividing bacteria, mycobacterial FtsK possesses an extended N-terminal region (**Fig. 1A**) that is conserved across both pathogenic and saprophytic mycobacterial species (**Fig. 1B**). This extension likely represents a mechanistic adaptation to asymmetric mode of division, mediating specific interaction with potential partners such as protein(s) and/or membrane lipid(s) to stabilize FtsK positioning and coordinate septal dynamics with chromosome translocation in the absence of canonical spatial regulators.

Removal of the N-terminal extension (nFtsK) results in a modest but reproducible decrease in cell length and a shorter generation time (**Fig. 2D**), without affecting septal recruitment, septum formation, or completion of division, indicating that the core divisiome remains functional. Our data support a model in which nFtsK stabilizes late-stage divisome dynamics and transiently delays cytokinesis (**Fig. 2D**). In its absence, the cell cycle progresses more rapidly but with reduced temporal precision. Although the average timing of chromosome segregation relative to FtsK disassembly is preserved, the variability of this coordination is significantly increased in the sFtsK strain. Thus, nFtsK does not alter the sequence of cell cycle events but constrains fluctuations around an optimal timing window, enhancing the robustness and fidelity of division rather than its speed.

The spatial organization of FtsK is strongly influenced by its N-terminal extension. sFtsK localizes closer to midcell than the full-length protein (**Fig. 2C**), indicating that the extension contributes to positioning FtsK at the non-central septa characteristic of mycobacteria. Despite increased confinement and reduced diffusion, sFtsK displays weaker clustering and broader spatial distributions at the division site. These findings indicate that confinement alone is insufficient for productive septal localization and suggest that nFtsK promotes selective confinement while suppressing off-target interactions. Loss of this spatial control likely contributes to the uncoupling of cell envelope biogenesis from DNA replication, as reflected by the increased variability in TMR-trehalose incorporation and replication initiation in daughter cells (**Fig. 2D**). Thus, the N-terminal extension appears relevant for coordinating septal synthesis with the replication cycle.

Changes in FtsK dynamics are accompanied by altered protein interactions (**Fig. 4B**, **Table 1**). Core divisome components associate with both full-length and truncated FtsK, consistent with preserved septal recruitment. However, sFtsK shows increased association with proteins linked to non-specific or aberrant interactions. Enrichment of FtsH in the sFtsK-HT pulldown suggests that, in the absence of nFtsK, FtsK may be more susceptible to quality-control surveillance, potentially due to inappropriate membrane contacts or altered interaction specificity. Full-length FtsK, by contrast, specifically associates with WhiA, a known division regulator, as well as enzymes involved in acetyl-CoA production and lipid biosynthesis, including Ino1. These associations suggest that nFtsK presumably couples late chromosome segregation to regulatory and metabolic processes required for septum maturation and new pole formation.

Our data further indicate that nFtsK acts primarily through protein–lipid interactions rather than direct DNA binding. Despite its strong positive charge, nFtsK does not bind DNA *in vitro* (**Fig. 4A**). Instead, it preferentially associates with cardiolipin and phosphatidic acid (**Fig. 4D**), lipids enriched at regions of negative membrane curvature such as the septum and cell poles. The ability of an isolated nFtsK-mNeonGreen fusion to localize to these regions, even in a heterologous system (**Fig. 4E**), demonstrates that this interaction is intrinsic and independent of other divisome components. Lipid-mediated targeting therefore provides a plausible mechanistic basis for how FtsK achieves robust spatial confinement in the absence of Min or nucleoid occlusion systems, while remaining sufficiently dynamic to support regulated assembly and disassembly.

In summary, the N-terminal extension of mycobacterial FtsK acts as a regulatory module that enhances the precision and robustness of late chromosome segregation during asymmetric cell division. Although dispensable for division under standard laboratory conditions, this domain constrains variability in the spatial and temporal execution of cytokinesis, a property likely important for fitness in fluctuating or stressful environments. More broadly, our findings illustrate how conserved bacterial cell-cycle proteins can evolve modular extensions to support non-canonical modes of growth and division.

## Material and Methods

### Plasmid Propagation and Bacteria Cultivation

Plasmids used for *Mycobacterium smegmatis mc^2^ 155* transformation were propagated in *Escherichia coli* DH5α. *E. coli* was cultured at 37°C in Luria-Bertani (LB) broth with shaking at 180 rpm or on LB agar plates (Difco). Media were supplemented with antibiotics (100 µg/ml ampicillin or 50 µg/ml kanamycin) and additives such as 0.004% X-Gal when necessary (Sambrook & Russell, 2001). *M. smegmatis* liquid cultures were grown at 37°C with agitation at 180 rpm in Middlebrook 7H9 broth supplemented with 0.05% Tween 80 and albumin-dextrose-catalase (ADC; BD), or in Difco Nutrient Broth (NB; BD). For solid media, *M. smegmatis* was grown on NB supplemented with 2% agar or on Middlebrook 7H10 agar with oleic acid-albumin-dextrose-catalase (OADC; BD). Cultures were incubated at 37°C for 2–5 days until visible colonies appeared. If required, media were supplemented with 100 µg/ml ampicillin, 50 µg/ml kanamycin, 0.004% X-Gal, 2% sucrose, or 1 mM IPTG.

### Bacterial staining and sample preparation

For snapshot analysis, *E. coli* BL21 (DE3) with either nFtsK-mNeonGreen or its negative control were grown overnight in LB liquid medium supplemented with chloramphenicol, and a small aliquot was used to inoculate fresh medium. Upon reaching an OD_600_ of 0.4–0.6, expression was induced by adding isopropyl-β-D-1-thiogalactopyranoside (IPTG) to a final concentration of 1 mM, followed by a 1-hour incubation. The cells where further centrifuged (5000 x g, 5min, RT), washed and resuspended in PBS.

For imaging, exponential-phase cultures of *M. smegmatis* grown in 7H9 supplemented with ADC and 0.05% Tween 80 (OD600 = 0.8–1.2) were harvested by centrifugation (5000 x g, 5 min, room temperature), washed once with PBS, and resuspended in PBS. Exponential-phase cells were used for staining. For cell membrane visualization, log-phase cells were stained with 0.5 µM FM5-95 dye (Thermo Fisher Scientific). For HaloTag labeling in *M. smegmatis* (FtsK-HT and sFtsK-HT), TMRDirect ligand (Promega, 100 µM stock) was added at a 1:1000 dilution to a final concentration of 100 nM. In all cases, 1 ml of the cell suspension was incubated with the respective dye for 30 minutes at 37°C with continuous agitation at 180 rpm. Following incubation, cells were centrifuged (5000 x g, 5 min, RT), washed, and resuspended in PBS.

### Fluorescence microscopy

Samples (*E. coli* or *M. smegmatis*) were mounted on Teflon-coated glass slides with 1.2% agarose. Snapshot analysis was performed using a Leica DM6 epifluorescence microscope equipped with an HC PL FLUOTAR 100x/1.32 OIL PH3 objective. The following imaging parameters were applied for FM5-95 (Thermo Fisher Scientific) and TMR Direct (Promega): Excitation 570–590 nm, emission 602–662 nm; exposure time 300 ms; excitation intensity 40%. For nFtsK-mNeonGreen: excitation 450–490 nm, emission 500–550 nm; exposure time 100 ms; excitation intensity 32%. Images were processed and analyzed using Fiji software and R software with the ggplot2 package.

### Time-lapse fluorescence microscopy (TLFM)

To visualize live cells, *Mycobacterium smegmatis* liquid cultures grown overnight (OD_600_ ≈ 0.5) were loaded into an ONIX microfluidic chamber. Specifically, 70 µl of the cell suspension was injected into the inlet of a B04A plate (Merck). Prior to loading, the initial two wells were primed with PBS, whereas the subsequent wells contained 150 - 300 µl of 7H9 medium enriched with ADC and 0.05% Tween80. After trapping the bacteria in the imaging chamber, a 45-minute wash was performed with fresh medium at a pressure of 3 psi, followed by continuous cultivation for 24 hours at 1.5 psi. For NADA-Green pulse chase labeling, the 2 minutes labelling was utilized every 60 minutes. Microscopy data were collected using phase contrast and fluorescence channels (475/28 nm excitation/525/48 nm emission for NADA-Green; 575/25 nm excitation/625/45 nm emission for HaloTag labeled with TMRDirect). The acquisition parameters included exposure times of 50–100 ms with 50% light intensity for the TMRDirect signal and 80–200 ms with 32% intensity for NADA-Green. Automated time-lapse imaging was conducted at 10-minute intervals on a DeltaVision Elite inverted microscope featuring a UPlanFL N 100×/1.3 Oil Ph3 objective and a temperature-controlled chamber set at 37 °C. Post-acquisition analysis was performed in Fiji (58), and data visualization was generated using the ggplot2 package in R (R Foundation for Statistical Computing, Austria).

### 3D-Structured Illumination Microscopy (3D-SIM)

3D-structured illumination microscopy (3D-SIM) imaging was conducted using a DeltaVision OMX SR system (GE Healthcare). The setup included 488 nm and 568 nm laser lines, a pco.edge 4.2 sCMOS camera, and an Olympus PlanApo 60x oil-immersion objective (1.42 NA). In 3D-SIM mode, Z-stacks were recorded as 20 optical sections with a 0.125 µm interval. Every section was generated using a striped illumination pattern across three rotation angles (−60°, 0°, +60°) and five phase shifts. FtsZ-EGFP and HupB-mCherry signals were acquired with 50% laser power and 20 ms exposure time. Image reconstruction and multi-channel alignment were executed using SoftWoRx software (GE Healthcare) applying default parameters.

### Single Particle Tracking (SPT)

Cells were cultured to mid-log phase in rich medium (7H9 supplemented with 10% ADC, 12.5 nM TMRDirect and 0.05% Tween 80). Slides and coverslips were prepared by overnight incubation in 1 M KOH, followed by a milli-Q water wash and drying with pressurized nitrogen. Prior to imaging, cells were spread onto agar pads (1% low melting agarose in 7H9 poured into 1.0 × 1.0 cm GeneFrames) and sealed with clean 0.17-mm coverslips. Imaging was performed using a Zeiss Elyra 7 inverted microscope equipped with an sCMOS 4.2 CL HS camera and an alpha Plan-Apochromat 63×/1.46 Oil objective. The Z-axis was maintained via the Definite Focus.2 system. Data were recorded at 37°C using a 20 ms exposure per frame over 10,000 frames. For (s)FtsK-HaloTag strain stained with TMRdirect, a 561 nm laser was used at 30% intensity in TIRF mode (62° angle) with 500mW power. For Single Particle Tracking (SPT) analysis, spots were detected and reconstructed into tracks using the TrackMate v6.0.1 plugin in Fiji. Spot identification utilized a 0.5 μm diameter with sub-pixel localization, a median filter, and a signal-to-noise threshold of 5. Track reconstruction for FtsK-HT and sFtsK-HT allowed a maximum linking distance of 0.7 μm with no frame gaps; only tracks exceeding four frames were included in the final analysis. Statistical processing and comparison of the trajectories were conducted in SMTracker 2.0. The movement of FtsK-HT and sFtsK-HT particles was characterized through dwell time, mean-squared displacement (MSD), and squared-displacement (SQD) analyses. Dwell time was calculated using a 100 nm confinement radius and fitted with two components. MSD values were derived from five time points and fitted to a linear equation. Finally, diffusion coefficients (D_xy_) in SQD analysis were determined from jump distance data (JD) using independent fitting for FtsK-HT and sFtsK-HT.

### Purification of His-nFtsK-FLAG

The obtained pellet of *E. coli* BL21 (DE3) pET28 nFtsK was resuspended in purification buffer A (PBS 10mM imidazole, supplemented with 50 μl of viscolase (A&A Biotechnology). The cell suspension was sonicated using a Sonics Vibracell sonicator for 15 minutes at 40% amplitude, with a pulse cycle of 5 seconds on and 5 seconds off at 4°C. The resulting lysate was transferred to 50 ml Falcon tubes and centrifuged in a Beckman Avanti centrifuge for 40 minutes at maximum speed. The collected supernatant was filtered through a 0.22 μm syringe filter. The filtered supernatant was loaded onto a GE Healthcare ÄKTA chromatography system using HisTrap nickel columns. Proteins specifically bound to the resin were eluted using 30 ml of linear gradient of elution buffer PBS 1.5M imidazole. Remaining protein bound to resin were eluted with HEPES 20mM, 500mM imidazole, 500mM NaCl, pH 7.2. The resulting protein preparations were analyzed by SDS-PAGE electrophoresis and Western Blot.

### Electrophoretic Mobility Shift Assay (EMSA)

The EMSA was performed using purified His-nFtsK-FLAG protein to analyze DNA-protein interactions. A constant concentration of DNA (50–100 nmol per sample) was incubated with increasing concentrations of nFtsK (0.15, 0.3, 0.6, 0.9, 1.25, and 1.5 μM) for 20 minutes at room temperature. Subsequently, the samples were resolved on a 1% agarose gel. Electrophoresis was carried out overnight at 4°C to maintain complex stability. The following day, the gel was stained in TBE buffer supplemented with ethidium bromide for 30 minutes. DNA bands were visualized using a UV transilluminator and captured with a UV-sensitive camera system.

### Lipid-Protein Interaction Assay

Lipid-protein interaction assays were performed using membrane lipid strips (Echelon Biosciences, P-6002) according to the manufacturer’s instructions. Briefly, 2 µl of a goat anti-mouse IgG secondary antibody conjugated to HRP (Invitrogen) and 2 µl of the His-nFtsK-FLAG protein were spotted onto the membrane as positive controls. The membrane was blocked overnight at 4°C with 5% non-fat dry milk in TBST (TBS + 0.1% Tween-20). The following day, the lipid strip was incubated for 1 h with 13 µg of His-nFtsK-FLAG protein in TBST buffer. After washing with TBST, the strip was incubated for 1 h with an anti-FLAG primary antibody (Sigma, F1804, 1:1000 dilution). Following further washes with TBST, the membrane was incubated with an HRP-conjugated secondary antibody (Invitrogen, 1:5000 dilution) for 1 h. Protein-lipid interactions were visualized using Pierce™ SuperSignal™ West Pico PLUS Chemiluminescent Substrate (Thermo Scientific) and detected with a ChemiDoc MP imaging system (Bio-Rad).

### Liquid chromatography–Mass spectrometry (LC-MS) sample preparation and analysis

LC-MS was performed on an M-Class Acquity UPLC connected to a Synapt XS HDMS equipped with a nanoESI source. Mobile phase A consisted of H2O + 0.1% FA, while mobile phase B of ACN + 0.1% FA. An 8–40% B 40min linear gradient at a 300nL/min flow rate was applied for sample separation on a C18 75μm x 250mm analytical column kept at 45°C. A 5-minute sample trapping step was performed prior to analytical column separation. Data were collected in ESI+ using ion mobility DDA (HDDDA), with MS scan rate of 0.6 seconds, and MS/MS scan rate of 0.2 seconds, both in the 50–2000 m/z range. The top 8 precursors from each MS scan with charges 2+, 3+, and 4+ were selected for MS/MS with 1 scan allowed per transition. Collision energy ramp determined specifically for each m/z (start LM, HM: 20–52 V; end LM, HM: 27–62 V) was applied on the transfer cell. Fragments not matching precursor’s drift time were stripped. Source and IMS conditions were fine-tuned. A (Glu1)-Fibrinopeptide B solution was acquired in-parallel as lockmass, and correction was applied post-acquisition. Four independent biological replicates were analyzed (n=4), 8 samples in total. Raw processing was performed using PLGS v3.0.3 (Waters). Obtained spectra were exported as .mgf files and searched using Mascot Server v2.8.0.1 (Matrix Science) against the Mycolicibacterium smegmatis protein sequence databank (UP000000757) to which porcine trypsin, human keratins, rat Ig gamma-2A C and Ig lambda-2 C sequences were appended (UniProt entries). Rat Ig sequences were identified in a SwissProt databank pre-search, and are most likely resin related. The search parameters were: peptide mass tolerance: 15 ppm; fragment mass tolerance: 25 ppm; max. protein mass: 1 MDa; digest enzyme: trypsin; max. missed cleavages: 3; variable modification: oxidation of methionine; FDR (PSM): 1%. Protein hits were grouped into families. The Mascot protein hit .csv files were exported for each search, and all families (322) were combined into a single list, along with their score sum, as well as peptide and occurrence counts for either Ftsk or sFtsk, separately. To distinguish proteins specific to FtsK or sFtsK, we assumed hits with no cooccurrence (hit must be present exclusively either in the FtsK pull-down or the sFtsK pull-down), and with a peptide count of at least two (minimal requirement hence being either 1 occurrence by 2 peptides or 2 occurrences with 1 peptide each). This way, 22 and 13 proteins were determined as exclusive for either FtsK or sFtsK, respectively. Data analysis can be found in SFxx, while further details are available in the repository files. The mass spectrometry proteomics data have been deposited to the ProteomeXchange Consortium via the PRIDE (Perez-Riverol et al., 2022) partner repository with the dataset identifier PXDPXD073126.

### Statistical Analysis and data visualization

Data were derived from three independent biological replicates for every investigated strain. The homogeneity of variance was verified using Levene’s test implemented in the *car* package within the RStudio environment (R Foundation for Statistical Computing, Austria). Based on the variance structure, statistical significance was determined using either the standard Student’s t-test or Welch’s t-test for unequal variances. Unless otherwise noted, numerical variations are reported as the standard deviation (SD), which is also represented by the error bars in the figures. Both statistical computations and graph generation were executed in R using the *stats* and *ggplot2* libraries. The thresholds for statistical significance were set as follows: *p* < *0.05* (\**), p < 0.01 (**), p < 0.001 (\*\*\**), and *p* < 0.0001 (****), while non-significant results are marked as ‘ns’. Schematics and model figures were designed using BioRender (BioRender.com).

## Acknowledgments

We thank Marta Kołodziej for providing preliminary data. The purchase of the M-Class Acquity-Synapt XS LC-MS system was supported by the “Excellence Initiative – Research University” program at the University of Wroclaw.

## Author contributions

J.Z.-C., J.H., K.M., contributed to the conception and design of the study. K.M., D.T., T.Ł., D.B. constructed strains. K.M., T.Ł., D.B. optimized and purified the proteins of interest. K.M. performed pulldown experiments, in silico analyses of FtsK sequences, microscopy experiments, and data analysis and visualization. M.T. performed mass spectrometry experiments and, together with K.M., analyzed the data. K.M. and J.Z.-C. wrote the original draft of the manuscript. J.H. reviewed and edited the manuscript. J.Z.-C. provided resources and secured funding. All authors contributed to the article and approved the submitted version.

## Materials & Correspondence

Materials and correspondence should be addressed to Jolanta Zakrzewska-Czerwińska.

## Data availability statement

All data supporting the findings of this study are included in the article and its supplementary materials. The mass spectrometry proteomics data have been deposited in the ProteomeXchange Consortium via the PRIDE partner repository under the dataset identifier PXD073126. Further inquiries can be directed to the corresponding author.

## References

Aldridge, B. B., Fernandez-Suarez, M., Heller, D., Ambravaneswaran, V., Irimia, D., Toner, M., & Fortune, S. M. (2012). Asymmetry and Aging of Mycobacterial Cells Lead to Variable Growth and Antibiotic Susceptibility. Science, 335(6064), 100–104. 10.1126/science.1216166

Bush, M. J., Casu, B., & Schlimpert, S. (2025). Dividing lines: compartmentalisation and division in Streptomyces. Current Opinion in Microbiology, 85, 102611. 10.1016/j.mib.2025.102611

Cameron, T. A., & Margolin, W. (2024). Insights into the assembly and regulation of the bacterial divisome. Nature Reviews Microbiology, 22(1), 33–45. 10.1038/s41579-023-00942-x

Chung, E. S., Kar, P., Kamkaew, M., Amir, A., & Aldridge, B. B. (2024). Single-cell imaging of the Mycobacterium tuberculosis cell cycle reveals linear and heterogenous growth. Nature Microbiology, 9(12), 3332–3344. 10.1038/s41564-024-01846-z

Crozat, E., & Grainge, I. (2010). FtsK DNA Translocase: The Fast Motor That Knows Where It’s Going. ChemBioChem, 11(16), 2232–2243. 10.1002/cbic.201000347

DeJesus, M. A., Gerrick, E. R., Xu, W., Park, S. W., Long, J. E., Boutte, C. C., Rubin, E. J., Schnappinger, D., Ehrt, S., Fortune, S. M., Sassetti, C. M., & Ioerger, T. R. (2017). Comprehensive Essentiality Analysis of the Mycobacterium tuberculosis Genome via Saturating Transposon Mutagenesis. MBio, 8(1). 10.1128/mBio.02133-16

Ginda, K., Santi, I., Bousbaine, D., Zakrzewska□Czerwińska, J., Jakimowicz, D., & McKinney, J. (2017). The studies of ParA and ParB dynamics reveal asymmetry of chromosome segregation in mycobacteria. Molecular Microbiology, 105(3), 453–468. 10.1111/mmi.13712

Grenga, L., Luzi, G., Paolozzi, L., & Ghelardini, P. (2008). The Escherichia coli FtsK functional domains involved in its interaction with its divisome protein partners. FEMS Microbiology Letters, 287(2), 163–167. 10.1111/j.1574-6968.2008.01317.x

Griffin, J. E., Gawronski, J. D., DeJesus, M. A., Ioerger, T. R., Akerley, B. J., & Sassetti, C. M. (2011). High-Resolution Phenotypic Profiling Defines Genes Essential for Mycobacterial Growth and Cholesterol Catabolism. PLoS Pathogens, 7(9), e1002251. 10.1371/journal.ppat.1002251

Jakimowicz, D., Brzostek, A., Rumijowska-Galewicz, A., Żydek, P., Dołzbłasz, A., Smulczyk-Krawczyszyn, A., Zimniak, T., Wojtasz, Ł., Zawilak-Pawlik, A., Kois, A., Dziadek, J., & Zakrzewska-Czerwińska, J. (2007). Characterization of the mycobacterial chromosome segregation protein ParB and identification of its target in Mycobacterium smegmatis. Microbiology, 153(12), 4050–4060. 10.1099/mic.0.2007/011619-0

Jean, N. L., Rutherford, T. J., & Löwe, J. (2020). FtsK in motion reveals its mechanism for double-stranded DNA translocation. Proceedings of the National Academy of Sciences, 117(25), 14202–14208. 10.1073/pnas.2001324117

Kieser, K. J., & Rubin, E. J. (2014). How sisters grow apart: mycobacterial growth and division. Nature Reviews Microbiology, 12(8), 550–562. 10.1038/nrmicro3299

Lutkenhaus, J. (2007). Assembly Dynamics of the Bacterial MinCDE System and Spatial Regulation of the Z Ring. Annual Review of Biochemistry, 76(1), 539–562. 10.1146/annurev.biochem.75.103004.142652

Mileykovskaya, E., Ryan, A. C., Mo, X., Lin, C.-C., Khalaf, K. I., Dowhan, W., & Garrett, T. A. (2009). Phosphatidic Acid and N-Acylphosphatidylethanolamine Form Membrane Domains in Escherichia coli Mutant Lacking Cardiolipin and Phosphatidylglycerol. Journal of Biological Chemistry, 284(5), 2990–3000. 10.1074/jbc.M805189200

Minato, Y., Gohl, D. M., Thiede, J. M., Chacón, J. M., Harcombe, W. R., Maruyama, F., & Baughn, A. D. (2019). Genomewide Assessment of Mycobacterium tuberculosis Conditionally Essential Metabolic Pathways. MSystems, 4(4). 10.1128/mSystems.00070-19

Renner, L. D., & Weibel, D. B. (2011). Cardiolipin microdomains localize to negatively curved regions of Escherichia coli membranes. Proceedings of the National Academy of Sciences, 108(15), 6264–6269. 10.1073/pnas.1015757108

Sassetti, C. M., Boyd, D. H., & Rubin, E. J. (2003). Genes required for mycobacterial growth defined by high density mutagenesis. Molecular Microbiology, 48(1), 77–84. 10.1046/j.1365-2958.2003.03425.x

Sivanathan, V., Emerson, J. E., Pages, C., Cornet, F., Sherratt, D. J., & Arciszewska, L. K. (2009). KOPS□guided DNA translocation by FtsK safeguards Escherichia coli chromosome segregation. Molecular Microbiology, 71(4), 1031–1042. 10.1111/j.1365-2958.2008.06586.x

Trojanowski, D., Hołówka, J., Ginda, K., Jakimowicz, D., & Zakrzewska-Czerwińska, J. (2017). Multifork chromosome replication in slow-growing bacteria. Scientific Reports, 7, 1–7. 10.1038/srep43836

Veiga, H., Jousselin, A., Schäper, S., Saraiva, B. M., Marques, L. B., Reed, P., Wilton, J., Pereira, P. M., Filipe, S. R., & Pinho, M. G. (2023). Cell division protein FtsK coordinates bacterial chromosome segregation and daughter cell separation in Staphylococcus aureus. The EMBO Journal, 42(11). 10.15252/embj.2022112140

Wu, K. J., Zhang, J., Baranowski, C., Leung, V., Rego, E. H., Morita, Y. S., Rubin, E. J., & Boutte, C. C. (2018). Characterization of conserved and novel septal factors in Mycobacterium smegmatis. Journal of Bacteriology, 200(6). 10.1128/JB.00649-17

Wu, L. J., & Errington, J. (2012). Nucleoid occlusion and bacterial cell division. Nature Reviews Microbiology, 10(1), 8–12. 10.1038/nrmicro2671

Yates, J., Zhekov, I., Baker, R., Eklund, B., Sherratt, D. J., & Arciszewska, L. K. (2006). Dissection of a functional interaction between the DNA translocase, FtsK, and the XerD recombinase. Molecular Microbiology, 59(6), 1754–1766. 10.1111/j.1365-2958.2005.05033.x

