## Supplementary figures and images for "FtsK with a unique N-terminal extension is involved in coordinating the final steps of chromosome segregation with asymmetrical division in mycobacterial cells"

### Figure S1

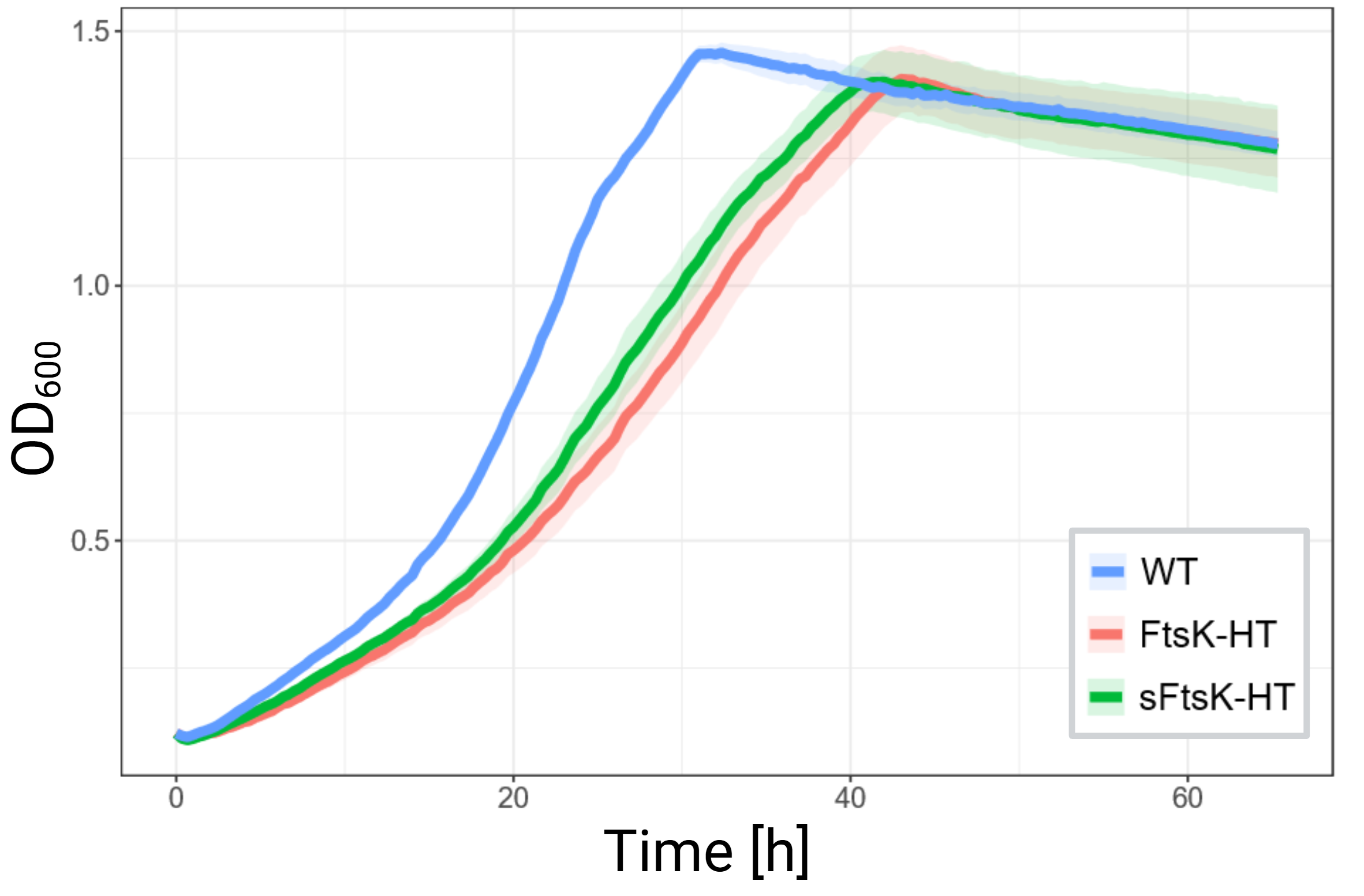

### Figure S2

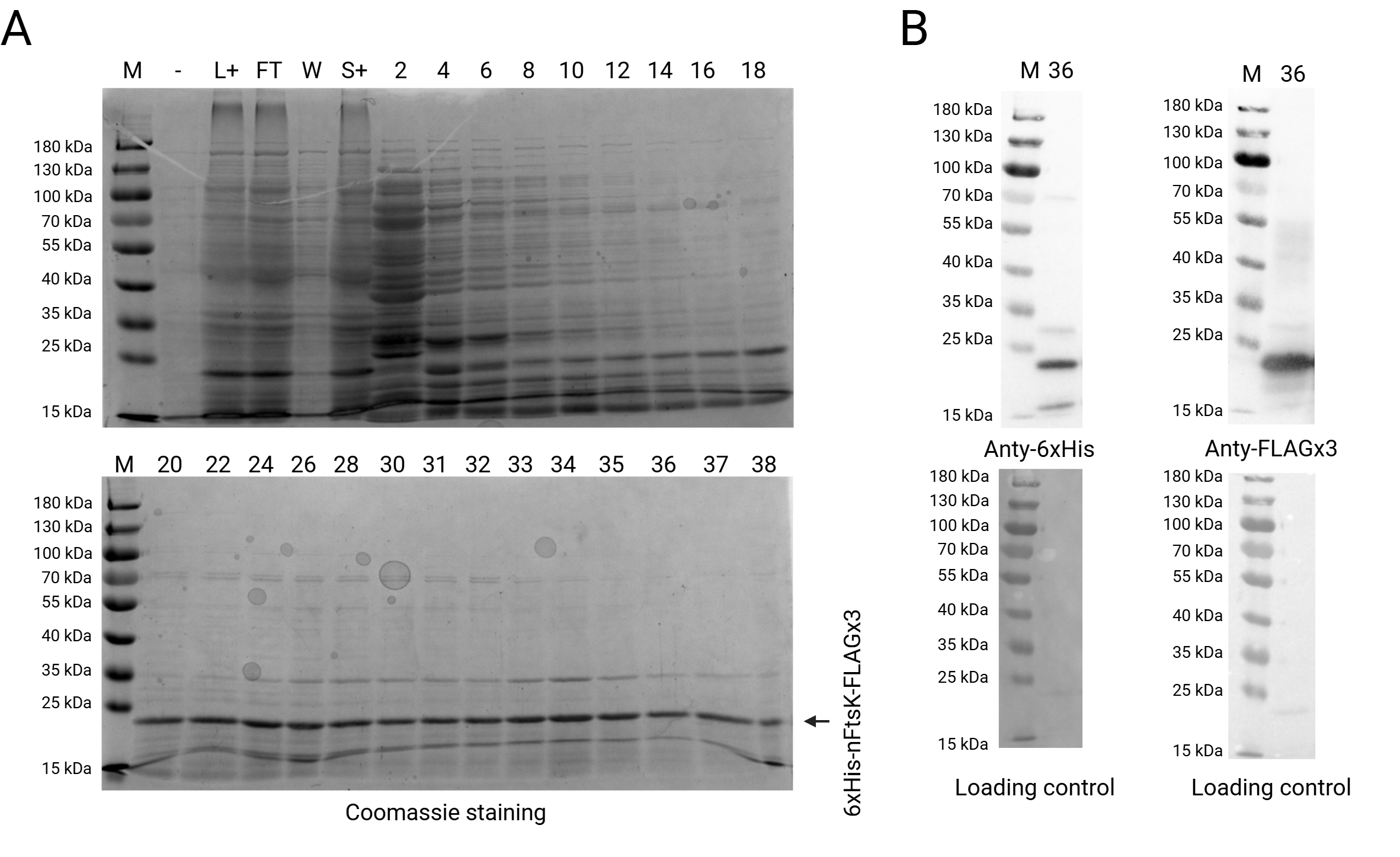
