## Supplementary material for "FtsK with a unique N-terminal extension is involved in coordinating the final steps of chromosome segregation with asymmetrical division in mycobacterial cells": Suplementary materials

### **Material and Methods**

**Construction of *M. smegmatis mc^2^* 155 mutant strains**

The allelic replacement of *ftsk* gene (*MSMEG_2690*) with fusion genes *ftsk-halotag or sftsk-halotag* was performed following the protocol outlined by Parish and Roberts (Parish & Roberts, 2015). Briefly, the chromosome of *M. smegmatis* *mc^2^ 155* was used as template to amplify region of *ftsk* gene using one primer set FtsK_p2NIL_FW x FtsK_p2NIL_RV for FtsK-HT strain and two primer sets for sFtsK_F1_p2NIL_FW x sFtsK_F1_p2NIL_RV and sFtsK_F2_p2NIL_FW x sFtsK_F2_p2NIL_RV. In the case of *halotag* fusion gene, it was amplified by PCR utilizing primer sets – FtsK_HT_p2NIL_FW x FtsK_HT_p2NIL_RV. Amplified products were cloned to p2NIL Ø plasmid and transformed colonies were spread on 7H10+OADC medium supplemented with kanamycin. Sequenced vectors were used in transformation of *M. smegmatis.*

**Construction *E. coli* BL21 (DE3) expressing nFtsK-mNeonGreen and microscopic analysis**

To visualize sFtsK in *E. coli* cells, PCR-amplified products were generated using two sets of primers: nFtsK_pACYC_Fw x link_nFtsK_pACYC_Rv and nFtsK_link_mNG_pACYC_Fw x pACYC_mNG_NheI_Rv (**Table 3**). The resulting fragments were cloned into the pACYCDuet™-1 vector (Sigma) via Sequence and Ligation Independent Cloning (SLIC). Beforehand, the vector was linearized using the restriction enzymes NcoI-HF and HindIII-HF (NEB). For the negative control, mNeonGreen was PCR-amplified using the primer set pACYC_KN_Link_mNG_F x pACYC_mNG_NheI_Rv. Transformants were selected on LB agar supplemented with chloramphenicol. The resulting plasmids were verified by PCR and sequencing, and then transformed into *E. coli* BL21(DE3) cells. Positive clones were selected for fluorescence microscopy experiments.

**SDS-PAGE and Western Blot**

Western blotting analysis was performed to assess the expression levels and purity of the fusion proteins. The eluted fractions were resolved by SDS-PAGE using 10% gels. Following electrophoresis, gels were either stained with InstantBlue® Coomassie Protein Stain (Abcam, ab119211) or processed for Western blotting by transferring the proteins onto a nitrocellulose membrane. To prevent non-specific binding, the membrane was blocked overnight at 4°C with 5% non-fat dry milk in TBST (TBS + 0.1% Tween-20). The primary antibodies utilized were anti-FLAG (Sigma, F1804, 1:1000 dilution) for His-nFtsK-FLAG and anti-6xHis (Invitrogen, MA1-135, 1:1000 dilution). Membranes were incubated with the primary antibody for 1 h at room temperature. After washing with TBST, the membranes were incubated for 1 h at room temperature with a goat anti-mouse IgG secondary antibody conjugated to HRP (Invitrogen, 1:5000 dilution). Protein bands were visualized using Pierce™ SuperSignal™ West Pico PLUS Chemiluminescent Substrate (Thermo Scientific) and captured with a ChemiDoc MP imaging system (Bio-Rad). Protein transfer and loading consistency were verified via Ponceau S staining (ThermoFisher Scientific) according to the manufacturer’s instructions.

**Pull-down assay**

Liquid cultures of strains expressing FtsK-HT and sFtsK-HT were cultivated to an optical density (OD_600_) of approximately 0.8. Biomass was harvested by centrifugation at 5,000 rpm for 20 minutes at 4°C, washed twice with ice-cold PBS, and resuspended in 10 mL of freshly prepared immunoprecipitation (IP) buffer (50 mM Tris-HCl pH 8.0, 250 mM NaCl, 0.8% Triton X-100) supplemented with a protease inhibitor cocktail (Pierce Protease Inhibitor, Thermo Scientific). Lysis was performed via sonication on ice for a total process time of 25 minutes (50% amplitude; 5-second pulse/5-second pause cycles). The resulting lysate was clarified by centrifugation at 10,000 rpm for 20 minutes at 4°C, followed by a second centrifugation step to ensure complete removal of cellular debris. Total protein concentration was quantified using the Bradford method (ROTI®Quant, Carl Roth). For the affinity purification, 10 mg of total protein lysate was diluted to a final volume of 14 mL with IP buffer. Magne HaloTag beads (100 µl; Promega) were equilibrated by washing three times in IP buffer to remove the storage solution, then added to the lysate and incubated overnight at 4°C with constant rotation. Following incubation, the beads were isolated using a magnetic rack and washed four times with 1 mL of cold IP buffer to remove non-specifically bound contaminants. To remove the detergent from the IP buffer, beads were washed four times in 25 mM Tris pH 7.5. Bead bound proteins were then denatured at 65°C for 10 minutes in 25 mM Tris pH 7.5, 0.1% sodium deoxycholate, 3mM DTT. Subsequently, 200 ng of trypsin was added to the sample for an overnight on-bead digestion in 37°C. Next day the solution was separated from the beads, acidified and sodium deoxycholate was removed by centrifugation. The supernatant was then desalted using a STAGE tip (Rappsilber et al., 2003). Obtained peptide pellet was resuspended in 0.1% formic acid (FA), 3% acetonitrile (ACN) solution.

### **Supplementary tables**

**Table S2. Primers used in the study**

| **Name** | **Sequence** | **Application** |
| --- | --- | --- |
| Ms_attB_L5_down | AGGCACATGCTGCCACTG | Strain construction |
| Ms_attB_L5_up | AGCGGATGCGCTACCAAG |  |
| sFtsK_F1_FW | GCATTAAAGCTTCACGTGGTCGACGTTCACCCGGATGCTGTGCGGG | Construction of *p2NIL sftsK* |
| sFtsK_F1_RV | GCGCGCGGCCGACCAAGGAGCCGTGCGTACAT |  |
| sFtsK_F2_FW | CACGGCTCCTTGGTCGGCCGCGCGCGT |  |
| sFtsK_F2_RV | GGGAATTCTTAATTAAGCGGCCGCGGTACCGATGGCGTCGGTCATCTGGTC |  |
| FtsK-HT_F1_FW | ATAAACTACCGCATTAAATCGTGGCGATCGTC | Construction of *p2NIL ftsK-halotag* |
| FtsK-HT_F1_RV | CGGTACCTTAACCGAACTCCTCGCCG |  |
| FtsK-HT_F2_FW | GGAGTTCGGTTAAGGTACCGGCTCGGC |  |
| FtsK-HT_F2_RV | CGGCAAGCTTAAGCTTTCAACCGGAAATCTCCAG |  |
| FtsK-HT_F3_FW | TTGAAAGCTTAAGCTTGCCGAGAGTCCTAC |  |
| FtsK-HT_F3_RV | TGACACTATAGAATACATAGGTGAACAGGAACACCAGGAAC |  |
| nFtsK_pACYC_Fw | TAACTTTAATAAGGAGATATACATGTTGCTCATACAACGATCACTGG | Construction of *pACYCDuet™-1 nftsK-mNeonGreen* |
| link_nFtsK_pACYC_Rv | AGCCGACATAAGCTTGTGCCCGGGCTCGAT |  |
| nFtsK_link_mNG_pACYC_Fw | ATCGAGCCCGGGCACAAGCTTATGTCGGCTGGCT |  |
| pACYC_mNG_NheI_Rv | TCGACTTAAGCATTATGCGGCCGCAGCTAGCTTATTTGTACAATTCATCCATGCC |  |
| pACYC_KN_Link_mNG_F | GTTTAACTTTAATAAGGAGATATACATGTCGGCTGGCTCCG | Construction of *pACYCDuet™-1 linker-mNeonGreen* |
| pACYC_mNG_NheI_Rv | TCGACTTAAGCATTATGCGGCCGCAGCTAGCTTATTTGTACAATTCATCCATGCC |  |
| FtsK_N-term_FW | GGTACCTTGCTCATACAACGATCACTGGAC | Construction *pET28 nftsk-FLAG* |
| FtsK_N-term_RV | CTCGAGGTGCCCGGGCTCGATGTCA |  |

**Table S3. Plasmids used in the study**

| **Name** | **Plasmid feature** | **Reference** |
| --- | --- | --- |
| p2NIL Ø | kanamycin resistance, *oriE*, suicide plasmid for allelic replacement | (Parish & Roberts, 2015) |
| pGOAL17 Ø | ampicillin resistance, *oriE*, selective PacI selective cassete with *lacZ*, *sacB* and *kanR* genes | (Parish & Roberts, 2015) |
| pMV_pAMI_ Ø | kanamycin resistance, *oriE*, inducible promotor *p_AMI_, attB* integrative plasmid for mycobacterial transformation | Lab collection |
| p2NIL ftsK-halotag | plasmid constructed on p2NIL Ø backbone with inserted *ftsk-halotag* | This study |
| p2NIL ftsK-halotag GOAL | plasmid constructed on *p2NIL ftsK-halotag* backbone with inserted *goal* cassette from pGOAL17 Ø. | This study |
| p2NIL sftsK | plasmid constructed on p2NIL Ø backbone with inserted DNA sequence of FtsK without first 141 aminoacids | This study |
| p2NIL sftsK GOAL | plasmid constructed on *p2NIL sftsK* backbone with inserted *goal* cassette from pGOAL17 Ø. | This study |
| pMVpNAT DnaN-mCherry | plasmid constructed on pMV_306_ Ø backbone with inserted promotor of DnaN and DnaN-mCherry | Lab collection |
| p2NIL HupB-mCherry GOAL | plasmid constructed on *p2NIL hupB-mCherry* backbone with inserted *goal* cassette from pGOAL17 Ø. | Hołówka et al., 2017 |
| pACYCDuet™-1 Ø | CmR, *cat* promoter, p15A *ori*, *lac* promotor, repressor, operator, His-tag, S-tag | Lab collection |
| pACYCDuet™-1 nftsK-mNeonGreen | plasmid constructed on pACYCDuet™-1 Ø backbone with inserted *nftsK-mNeonGreen* | This study |
| pACYCDuet™-1 linker-mNeonGreen | plasmid constructed on pACYCDuet™-1 Ø backbone with inserted *linker-mNeonGreen* | This study |
| pET28a(+) Ø | kanamycin resistance, pBR322 origin, T7 promoter/expression system with N-terminal His-tag, thrombin cleavage site, and T7 tag. 3xFLAG sequence inserted af | Lab collection |
| pET28a nftsK-FLAGx3 | plasmid constructed on pET28a(+) Ø backbone with inserted *nftK-FLAGx3* | This study |

**Table S4. Strains used in the study**

| **Name** | **Genotype** | **Source** |
| --- | --- | --- |
| WT | *M. smegmatis mc^2^ 155* | Lab collection |
| FtsK-HT | *M. smegmatis mc^2^ 155 ftsK-halotag* | This study |
| sFtsK-HT | *M. smegmatis mc^2^ 155 sftsK-halotag* | This study |
| FtsK-EGFP HupB-mCherry | *M. smegmatis mc^2^ 155 ftsK-egfp hupB-mCherry* | This study |
| FtsK-HT DnaN-mCherry | *M. smegmatis mc^2^ 155 ftsK-halotag attBL5::pMV306_pnat dnaN-mCherry_* | This study |
| sFtsK-HT DnaN-mCherry | *M. smegmatis mc^2^ 155 sftsK-halotag attBL5::pMV306_pnat dnaN-mCherry_* | This study |
| FtsK-HT HupB-mCherry | *M. smegmatis mc^2^ 155 ftsK-halotag attBL5::pMV306_pnat hupB-mCherry_* | This study |
| sFtsK-HT HupB-mCherry | *M. smegmatis mc^2^ 155 sftsK-halotag attBL5::pMV306_pnat hupB-mCherry_* | This study |
| His-nFtsK-FLAG | *E. coli BL21 (DE3) pET28 His-nFtsK-FLAG* | This study |
| KN-mNG | *E. coli BL21 (DE3) pACYC linker-mNeonGreen* | This study |
| nFtsK-mNG | *E. coli BL21 (DE3) pACYC nFtsK-mNeonGreen* | This study |

### **Supplementary figures**

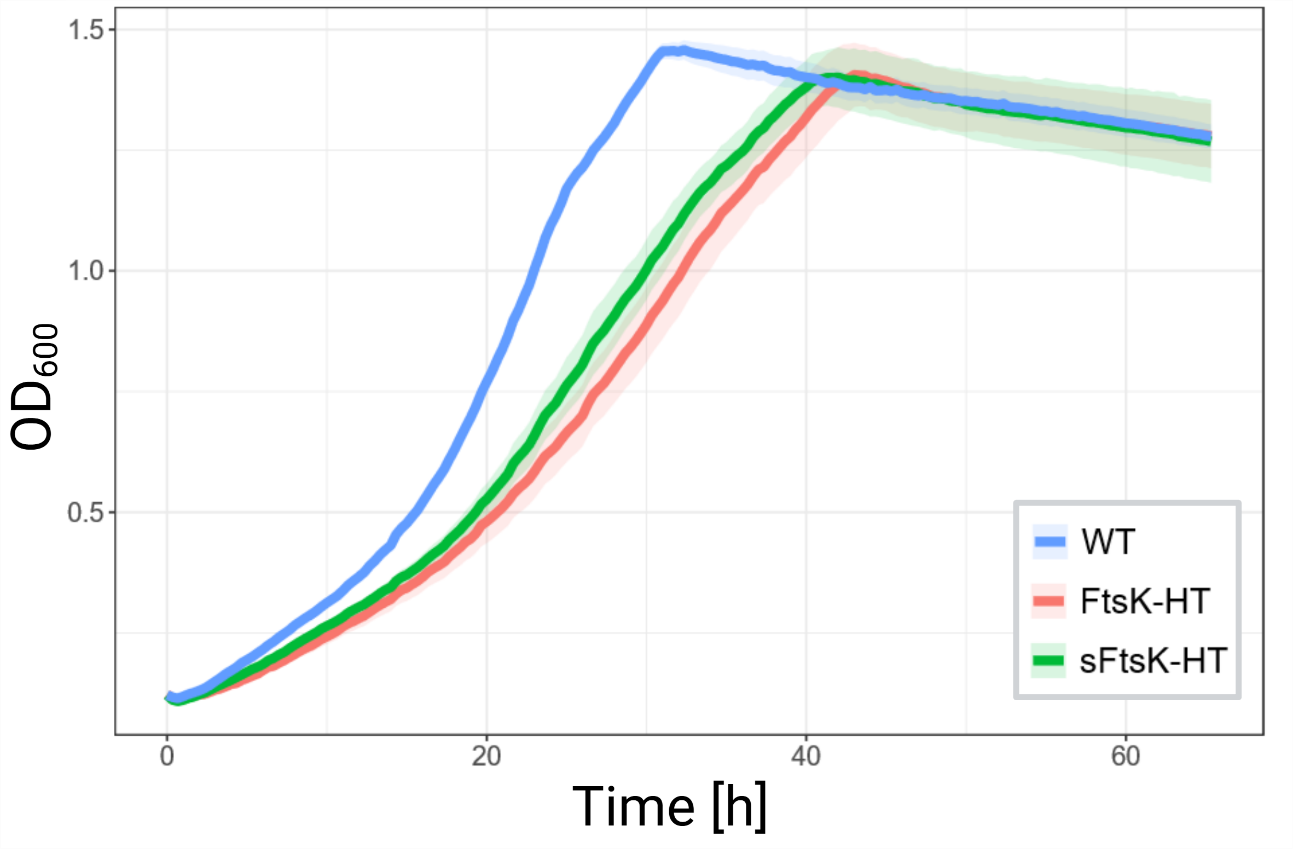

**Fig. S1. Growth rates of *M. smegmatis* strains.** Curves represent the mean optical density measured at 600 nm (OD_600_) over time. Shaded areas represent the standard deviation (n=6). Both strains expressing sFtsK-HT show slightly reduced growth in comparison to the wild type, while showing no significant differences between each other.

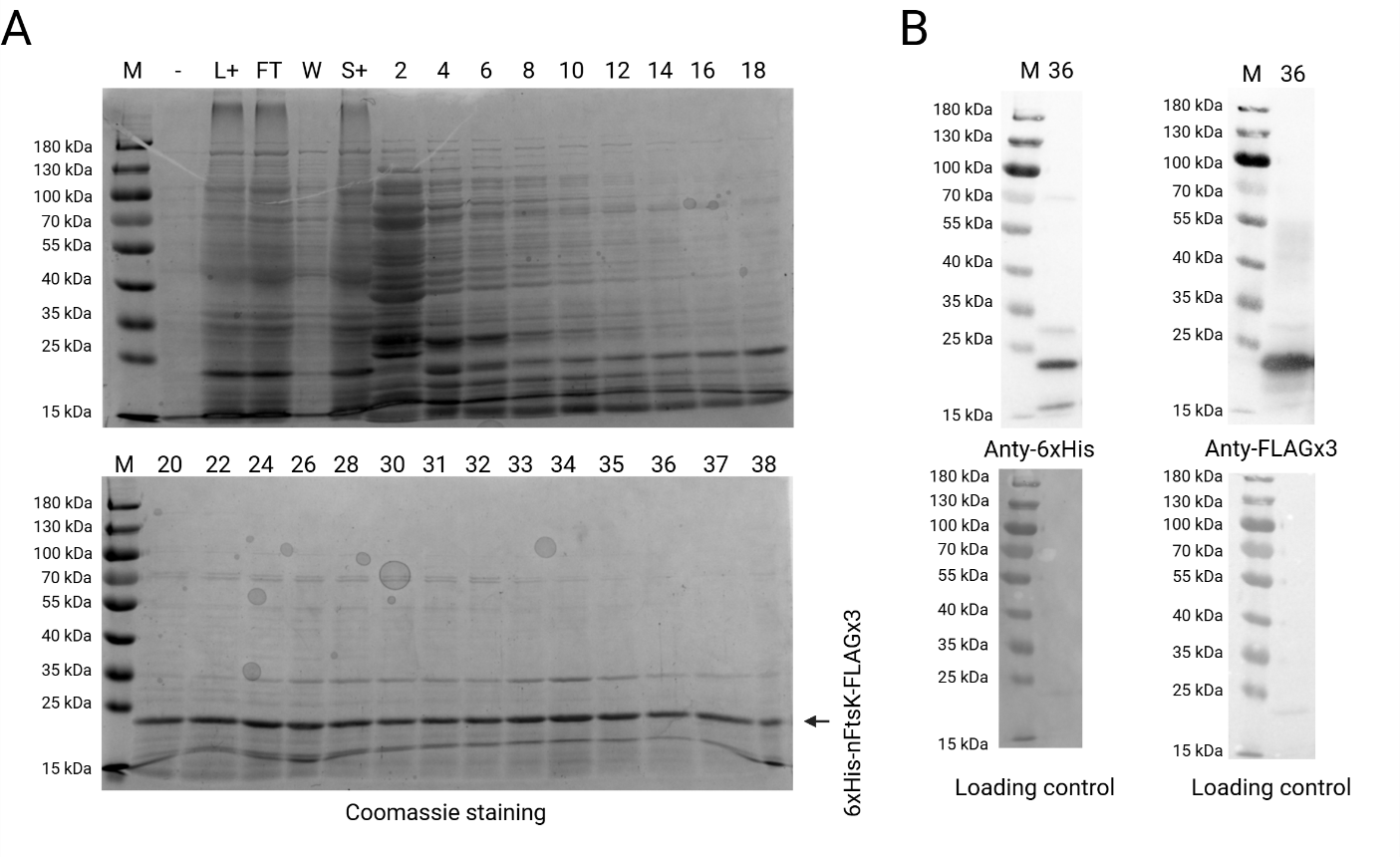
**Figure S2. Purification and quality assessment of 6xHis-nFtsK-FLAG. A.** SDS-PAGE analysis of eluted fractions from 6xHis-nFtsK-FLAG purification. Fraction 36 was selected for subsequent downstream analysis. The predicted molecular weight of 6xHis-nFtsK-FLAGx3 is 22.6 kDa. **B.** Western blot analysis of fraction 36 to confirm the identity of the target protein and determine whether additional bands represent degradation products or other species of the target protein. The primary band corresponds to the estimated molecular weight of 6xHis-nFtsK-FLAGx3 (22.6 kDa).

Parish, T., & Roberts, D. M. (2015). Mycobacteria protocols: Third edition. In T. Parish & D. M. Roberts (Eds.), *Mycobacteria Protocols: Third Edition* (Vol. 1285). Springer New York. https://doi.org/10.1007/978-1-4939-2450-9

Rappsilber, J., Ishihama, Y., & Mann, M. (2003). Stop and Go Extraction Tips for Matrix-Assisted Laser Desorption/Ionization, Nanoelectrospray, and LC/MS Sample Pretreatment in Proteomics. *Analytical Chemistry*, *75*(3), 663–670. https://doi.org/10.1021/ac026117i
